# Complete mitochondrial genome assemblies for three species of searobins (genera *Bellator* and *Prionotus*) and their phylogenetic relationships within Triglidae: Prionotinae

**DOI:** 10.1101/2025.02.01.635949

**Authors:** Alan Marín, Ruben Alfaro, Eliana Zelada-Mázmela

**Affiliations:** Laboratorio de Genética, Fisiología y Reproducción, Facultad de Ciencias, Universidad Nacional del Santa, Chimbote, Perú; Ecobiotech Lab S.A.C. Trujillo, Perú

**Keywords:** Mitogenomics, Phylogeny, Scorpaenoidei, GenBank data mining

## Abstract

Species from the genera *Bellator* and *Prionotus*, commonly known as ‘searobins’, are marine ray-finned fish that belong to the subfamily Prionotinae. These fish have evolved distinctive morphological features and specialized behaviors that enable them to walk along the seafloor while detecting buried prey. This unique adaptation makes them ideal candidates for studies in evolutionary genetics. However, their phylogenetic relationships remain poorly understood. In this study, we utilized publicly available genomic reads from the GenBank database to assemble the first complete mitochondrial genomes of three searobin species: *Bellator militaris, Prionotus alatus*, and *P. stephanophrys*. We conducted a comparative analysis of these mitochondrial genomes, including the first phylogenetic analysis of the Prionotinae based on complete mitogenomic data. The resulting circular contigs measured 16,765 base pairs for *B. militaris*, 16,602 base pairs for *P. alatus*, and 16,896 base pairs for *P. stephanophrys*. The three mitogenomes exhibited a typical vertebrate organization, which includes 13 protein-coding genes, 2 ribosomal RNAs, 22 transfer RNAs, and a putative control region. Notably, *P. stephanophrys* contained an additional tRNA-Leu and an extra non-coding region. A Bayesian phylogenetic analysis grouped all *Bellator* species into a monophyletic clade within Prionotinae, while the sister taxon *Prionotus* formed a separate monophyletic subclade. The findings of this research provide valuable insights into the phylogenetic relationships and evolutionary history of the genera *Bellator* and *Prionotus*. Clarifying the taxonomy of these species may also support future management and conservation efforts for these economically valuable species.

## 1. Introduction

Species from the genera *Bellator* Jordan & Evermann, 1896 and *Prionotus* Lacepède, 1801 (Triglidae), commonly known as ‘searobins’, are the only accepted sister taxa within the subfamily Prionotinae (Portnoy et al. 2017). These species not only support important fisheries across their distribution ranges but also exhibit specialized morphological and biological characteristics. One notable feature is their wing-like pectoral fins, which have three modified lower rays that allow them to ‘walk’ along the sea bottom (Portnoy et al. 2017). In some species, these modified rays are sensitive to chemical and mechanical stimuli, enabling them to detect buried prey (Portnoy et al. 2017; Allard et al. 2024). These unique features make searobins ideal biological models for studies in evolutionary genetics and adaptation (Herbert et al. 2024). Prionotinae species are exclusively found in the Americas (Portnoy et al. 2017). According to the World Register of Marine Species (WoRMS) database (available at https://www.marinespecies.org/index.php), there are currently eight extant *Bellator* species— four from the eastern Pacific and four from the western Atlantic. Additionally, there are 23 extant *Prionotus* species, with 16 located in the western Atlantic and seven in the eastern Pacific.

Taxonomic research on the classification of these species is still limited. Only a few studies have investigated their taxonomy using morphological traits (Imamura, 1996; Richards and Jones, 2002). Additionally, only one study has conducted a molecular phylogenetic analysis based on partial mitochondrial and nuclear gene fragments (Portnoy et al. 2017). While there has been some consistency between the findings of previous morphological studies and molecular phylogenetic analyses, the molecular research conducted by Portnoy et al. (2017) suggests that the subfamily Prionotinae has a monophyletic origin, with the genus *Bellator* nested within a paraphyletic *Prionotus*. The authors concluded that the currently recognized genus *Bellator* should be regarded as a junior synonym of *Prionotus*. These findings highlight the need for further studies using robust phylogenetic methods and encompassing a broader species coverage to better clarify their taxonomic status.

Due to the limitations of phylogenetic relationships and limited taxonomic resolution derived from morphological characters or partial gene fragments, there is and increasing need for more comprehensive genome-wide data to address complex phylogenetic issues within fish lineages (Miya and Nishida, 2015). In this regard, nucleotide information extracted from complete mitochondrial genomes (mitogenomes) has been effective in clarifying the phylogeny of various fish taxa across different taxonomic levels (Miya and Nishida, 2015). Moreover, the unique characteristics of mitogenomes—such as their fast evolutionary rates, absence of recombination, and high copy numbers—make them an excellent source of powerful molecular markers. These markers are suitable for diverse applications, including accurate species identification, wildlife forensics, environmental DNA analysis, population genetics, and evolutionary studies (Maduna et al. 2022).

Despite the availability of extensive genomic data from next-generation sequencing (NGS) in the GenBank database (Eldem and Balcı, 2024), there are only a few complete mitogenomes from Prionotinae species, and no studies have conducted a phylogenetic analysis based on mitogenomic information to investigate their evolutionary relationships. The objectives of the present work are: a) to assemble the mitochondrial genomes of commercially important searobin species using Sequence Read Archive (SRA) data mined from GenBank; b) to describe the characteristics of these mitogenomes and perform comparative analyses; and c) to investigate the phylogenetic relationships among Triglidae species by analyzing the assembled Prionotinae mitogenomes in conjunction with all available Triglidae mitogenomes in the GenBank database.

## 2. Materials and methods

### 2.1 Genomic NGS reads mining, mitogenome assembly and annotation

We mined short genomic read sequences (SRS) from publicly available SRA repository data available in the GenBank database. Table 1 presents the metadata for the three searobin specimens including the collection site and date, BioSample and experiment accessions, and the sequencing technology platform. It also outlines the main characteristics of the short read-sequences (pre and post processing) obtained from GenBank. The three searobin specimens were collected by the Smithsonian National Museum of Natural History as part of their project entitled ‘NOAA Genome Skimming of Marine animals inhabiting the US Exclusive Economic Zone’ (BioProject accession PRJNA720393). They consist of *B. militaris* (BioSample SAMN36762063), *P. alatus* (BioSample SAMN36762230), and *P. stephanophrys* (BioSample SAMN31811746).

**Table 1.** Metadata information of the three searobin species, genomic read sequences retrieved from the GenBank database, and mitochondrial genome assembly results.

|  | SPECIES |  |  |
| --- | --- | --- | --- |
|  | *Bellator militaris* | *Prionotus alatus* | *Prionotus stephanophrys* |
| **METADATA** |  |  |  |
| BioSample | SAMN36762063 | SAMN36762230 | SAMN31811746 |
| Submitted by | Smithsonian National Museum of Natural History | Smithsonian National Museum of Natural History | Smithsonian National Museum of Natural History |
| BioProject | PRJNA720393 | PRJNA720393 | PRJNA720393 |
| Collection site | Atlantic Ocean, USA | Atlantic Ocean, USA | Pacific Ocean in El Salvador |
| Collection date | October 18, 2022 | September 17, 2022 | December 10, 2010 |
| Coordinates | 34.9233 N, 75.4535 W | 34.6094 N, 75.7052 W | 12.7459 N, 89.0088 W |
| NGS Platform | Illumina NovaSeq 6000 | Illumina NovaSeq 6000 | Illumina NovaSeq 6000 |
| Strategy | WGS | WGS | WGS |
| SRA | SRR25465203 | SRR25465369 | SRR22396965 |
| Gbp | 1.9 | 1.9 | 1.8 |
| Reads (millions) | 12.8 | 12.5 | 12.1 |
| **ASSEMBLY DATA** |  |  |  |
| Post trimming (millions) | 12.7 | 12.49 | 12.07 |
| Mapped to Ref Seq | *B. brachychir* (OP056895) | *P. nudigula* (NC_088019) | *P. evolans* (NC_083024) |
| Assembled reads | 395,513 | 30,235 | 24,089 |
| **RESULTS** |  |  |  |
| GenBank accession | BK070174 | BK069871 | BK071384 |
| Submission date | 19 February 2025 | 25 January 2025 | 24 April 2025 |
| Mito length bp | 16,765 | 16,602 | 16,896 |
| A % | 26.9 | 26.3 | 26.6 |
| C % | 29.3 | 29.6 | 29.6 |
| G % | 17.6 | 17.3 | 17.6 |
| T % | 26.3 | 26.8 | 26.2 |

The raw paired-end FASTQ genomic reads underwent quality control using FastQC and were trimmed with BBDuk, as implemented in Geneious Prime v. 2025.0 (Biomatters Ltd., Auckland, New Zealand). To validate the mitogenomes assembled herein, we used two different approaches (‘map to reference’ and a hybrid ‘map to reference/de novo’ approach) following Alfaro et al. (2025). The reference sequences information used in the ‘map to reference’ approach is shown in Table 1.

For the annotation of the mitochondrial genes, we used the ‘Annotate and Predict’ feature of Geneious Prime by comparing the novel *B. militaris, P. alatus*, and *P. stephanophrys* mitogenomes to the complete mitogenomes of closely related species. The start and stop codon positions of the protein-coding genes (PCGs) were confirmed using the Open Reading Frame Finder tool (https://www.ncbi.nlm.nih.gov/orffinder/) using the vertebrate mitochondrial genetic code. The positions of the transfer RNA genes (tRNAs) were corroborated using the tRNAscan-SE tool (Chan and Lowe, 2019). The circular mitogenome maps were produced with Organellar Genome Draw ‘OGDRAW’ version 1.3.1 (Greiner et al. 2019) and then subjected to manual edition with Inkscape macOS version 1.4 (Inkscape Project, 2020).

### 2.2 Bayesian phylogenetic analysis

For the Bayesian phylogenetic analysis, we used the 13 PCGs from the three mitogenomes assembled in this work along with those from all Triglidae species available in GenBank, totaling 20 species (11 from Prionotinae and 9 from Triglinae). A Hoplichthyidae species, *Hoplichthys gilberti*, was used as outgroup. The nucleotide sequences from each PCG obtained from each mitogenome were multialigned using MEGA 7 (Kumar et al. 2016) and concatenated with SeaView version 5.0.4 (Guoy et al. 2021). A final matrix of 11,415 nucleotides was obtained and used in subsequent phylogenetic analysis. The program jModelTest 2 (Darriba et al. 2012) was used to find the most appropriate model of nucleotide evolution for each one of the 13 PCGs under the corrected Bayesian Information Criterion (BICc). To assess the levels of substitution saturation at single codon positions from each PCG we used Xia’s method implemented in DAMBE7 (Xia, 2018). As a result, we observed that the third codon positions of three PCGs (ATP8, ND3, and ND6) showed signs of saturation, prompting us to conduct the Bayesian analysis with data partitioned by gene and codon position excluding the third codon position from those genes. The Bayesian phylogenetic trees were constructed with MrBayes 3.2.7 (Ronquist et al. 2012) using the CIPRES Science Gateway 3.3 server (Miller et al. 2010). Two independent runs of four Markov chains each were executed for 10,000,000 generations, with a sampling frequency of 1,000. We discarded the first 25% of the sampled trees as burn-in. The final consensus phylogenetic tree was visualized using Figtree version 1.4.4 (http://tree.bio.ed.ac.uk/software/figtree/).

## 3. Results and discussion

### 3.1 Mitochondrial genome assemblies

In this work, we assembled and annotated the first mitochondrial genomes of three important searobin species: the hornet searobin *B. militaris* (found from North Carolina through the Gulf of Mexico), the spiny searobin *P. alatus* (occurring from Virginia in the USA to the Campeche Shelf in Mexico), and the lumptail searobin *P. stephanophrys* (found from the Columbia River in Washington in the USA to Ilo in Peru) (Russell et al. 1992; Chirichigno and Cornejo, 2001). There were no differences between the consensus mitogenome contigs obtained from the two different assembly approaches applied herein. The number of total assembled reads and the base frequencies for each mitogenome are shown in Table 1. The mitogenome of *B. militaris* consisted of a single circular contig of 16,765 (GenBank accession BK070174), for *P. alatus* it was 16,602 bp (GenBank accession BK069871), and for *P. stephanophrys* it was 16,896 bp (GenBank accession BK071384) (Table 1, Fig. 1). These lengths fall within the range reported of for other *Bellator* and *Prionotus* mitogenomes, which vary from 16,599 bp in *P. nudigula* (NC_088019) to 16,947 in *B. egretta* (NC_083033).

**Fig. 1.**
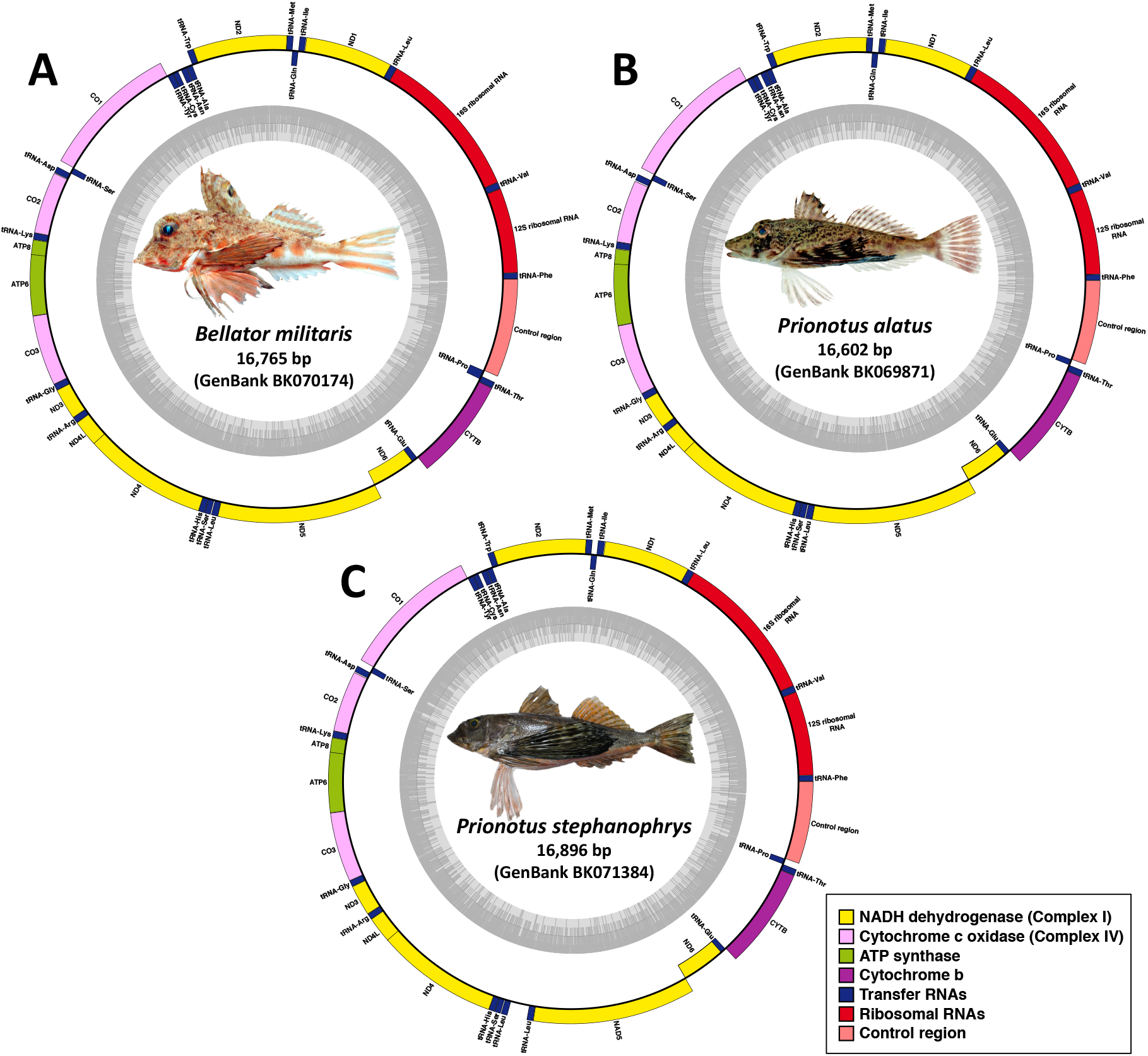
Circular mitochondrial genome organization of A) *Bellator militaris*, B) *Prionotus alatus* (specimen photographs kindly provided by R. Robertson; Robertson and van Tassell, 2015), and C) *P. stephanophrys* (photo by R. Alfaro). Genes located in inner circle (negative strand) are transcribed in a clockwise direction, while those on the outer circle (positive strand) are transcribed counterclockwise. The innermost circle of the GC content graph indicates a 50% threshold, with darker and lighter grey shades representing GC and AT content, respectively. The different gene types are illustrated with colored bars: ND dehydrogenase (yellow), cytochrome c oxidase (pink), ATP synthase (green), cytochrome b (purple), transfer RNA (blue), and ribosomal RNA (red). The control region is represented by a coral red bar.

### 3.2 Mitochondrial genome characterization

The mitogenomes showed a typical vertebrate organization, consisting of 13 PCGs, 2 ribosomal RNAs, 22 transfer RNAs, and a putative control region (Table 2). It is worth noting that the mitogenome of *P. stephanophrys* displayed an additional tRNA-Leu and an extra non-coding region. This extra tRNA-Leu was found on the positive strand, situated between a 220 bp non-coding region located downstream of the conserved block ND4/His/Ser/Leu and the ND5 gene (Table 1 and Fig. 1). A similar non-coding region was also identified in the mitogenomes of *P. rubio* (219 bp, Genbank OP056884), *P. tribulus* (219 bp, Genbank OP056925), and *P. evolans* (235 bp, Genbank NC_080324), which are closely related to *P. stephanophrys* (Fig. 2). Similarly, non-coding regions of 533 bp and 165 bp were reported within the mitogenome cluster ND4/His/Ser/Leu/ND5 in the freshwater gobioid *Odontobutis platycephala* (GenBank accession NC_010199; Ki et al. 2008) and in the threadfin *Eleutheronema rhadinum* (Genbank accession MW845829; Zhong et al. 2021) respectively.

**Table 2.**
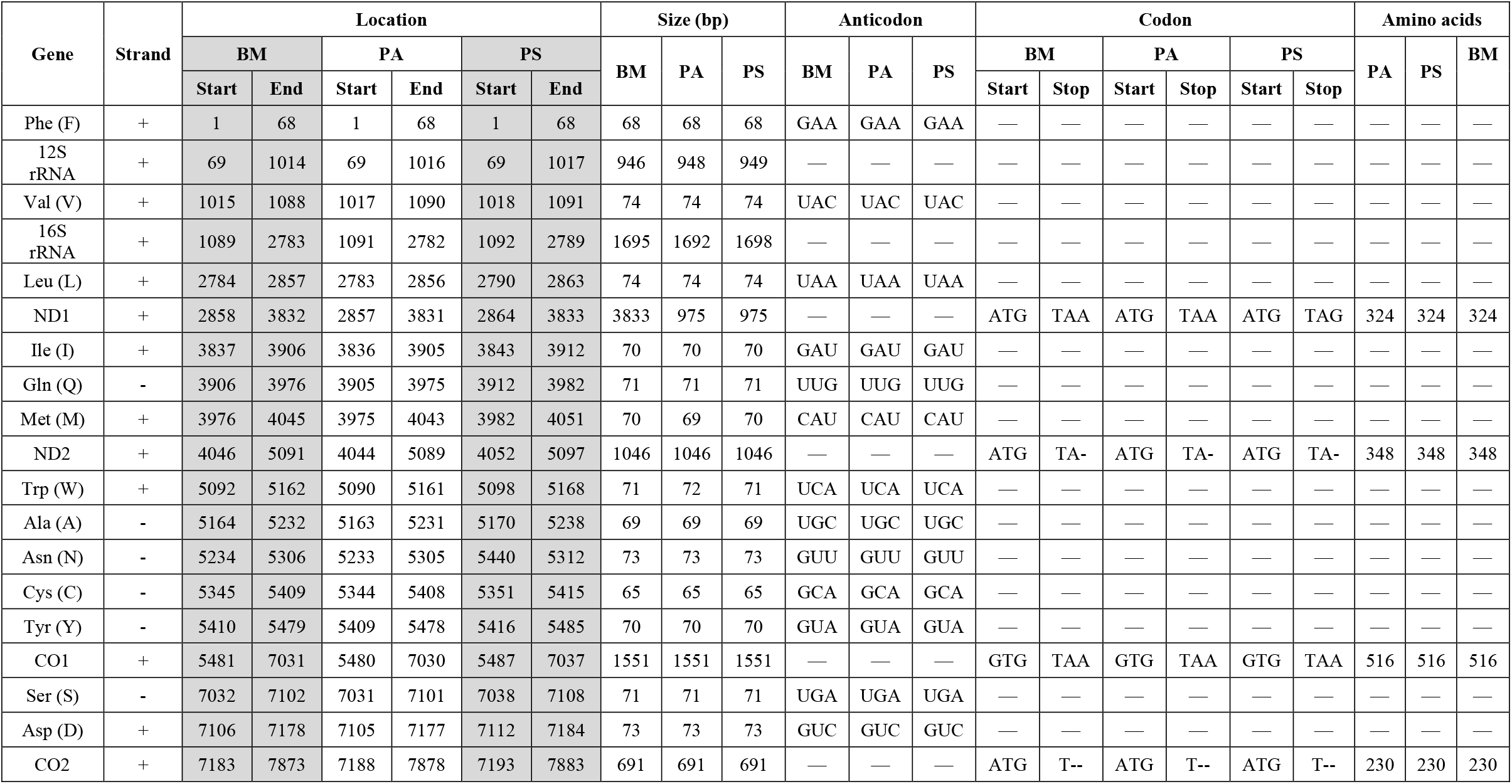
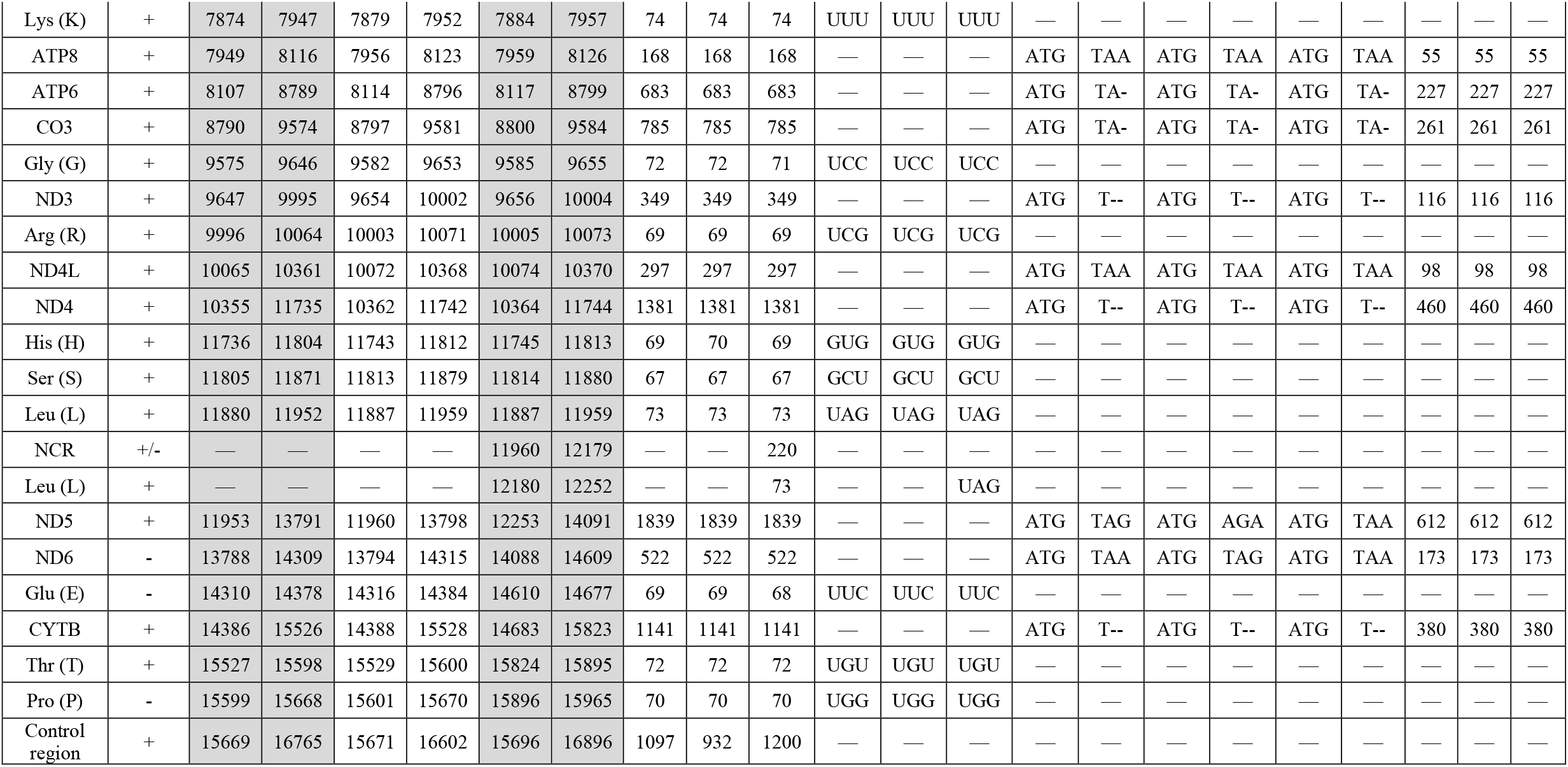
Complete mitochondrial annotation and number of amino acids for each gene identified in the mitochondrial genomes of *Bellator militaris* (BM), *Prionotus alatus* (PA), and *P. stephanophrys* (PS). The plus ‘+’ and minus ‘-’ symbols represent the heavy and light strands, respectively. NCR: non-coding region.

| Gene | Strand | Location |  |  |  |  |  | Size (bp) |  |  | Anticodon |  |  | Codon |  |  |  |  |  | Amino acids |  |  |
| --- | --- | --- | --- | --- | --- | --- | --- | --- | --- | --- | --- | --- | --- | --- | --- | --- | --- | --- | --- | --- | --- | --- |
|  |  | BM |  | PA |  | PS |  | BM | PA | PS | BM | PA | PS | BM |  | PA |  | PS |  | PA | PS | BM |
|  |  | Start | End | Start | End | Start | End |  |  |  |  |  |  | Start | Stop | Start | Stop | Start | Stop |  |  |  |
| Phe (F) | + | 1 | 68 | 1 | 68 | 1 | 68 | 68 | 68 | 68 | GAA | GAA | GAA | — | — | — | — | — | — | — | — | — |
| 12S rRNA | + | 69 | 1014 | 69 | 1016 | 69 | 1017 | 946 | 948 | 949 | — | — | — | — | — | — | — | — | — | — | — | — |
| Val (V) | + | 1015 | 1088 | 1017 | 1090 | 1018 | 1091 | 74 | 74 | 74 | UAC | UAC | UAC | — | — | — | — | — | — | — | — | — |
| 16S rRNA | + | 1089 | 2783 | 1091 | 2782 | 1092 | 2789 | 1695 | 1692 | 1698 | — | — | — | — | — | — | — | — | — | — | — | — |
| Leu (L) | + | 2784 | 2857 | 2783 | 2856 | 2790 | 2863 | 74 | 74 | 74 | UAA | UAA | UAA | — | — | — | — | — | — | — | — | — |
| ND1 | + | 2858 | 3832 | 2857 | 3831 | 2864 | 3833 | 3833 | 975 | 975 | — | — | — | ATG | TAA | ATG | TAA | ATG | TAG | 324 | 324 | 324 |
| Ile (I) | + | 3837 | 3906 | 3836 | 3905 | 3843 | 3912 | 70 | 70 | 70 | GAU | GAU | GAU | — | — | — | — | — | — | — | — | — |
| Gln (Q) | - | 3906 | 3976 | 3905 | 3975 | 3912 | 3982 | 71 | 71 | 71 | UUG | UUG | UUG | — | — | — | — | — | — | — | — | — |
| Met (M) | + | 3976 | 4045 | 3975 | 4043 | 3982 | 4051 | 70 | 69 | 70 | CAU | CAU | CAU | — | — | — | — | — | — | — | — | — |
| ND2 | + | 4046 | 5091 | 4044 | 5089 | 4052 | 5097 | 1046 | 1046 | 1046 | — | — | — | ATG | TA- | ATG | TA- | ATG | TA- | 348 | 348 | 348 |
| Trp (W) | + | 5092 | 5162 | 5090 | 5161 | 5098 | 5168 | 71 | 72 | 71 | UCA | UCA | UCA | — | — | — | — | — | — | — | — | — |
| Ala (A) | - | 5164 | 5232 | 5163 | 5231 | 5170 | 5238 | 69 | 69 | 69 | UGC | UGC | UGC | — | — | — | — | — | — | — | — | — |
| Asn (N) | - | 5234 | 5306 | 5233 | 5305 | 5440 | 5312 | 73 | 73 | 73 | GUU | GUU | GUU | — | — | — | — | — | — | — | — | — |
| Cys (C) | - | 5345 | 5409 | 5344 | 5408 | 5351 | 5415 | 65 | 65 | 65 | GCA | GCA | GCA | — | — | — | — | — | — | — | — | — |
| Tyr (Y) | - | 5410 | 5479 | 5409 | 5478 | 5416 | 5485 | 70 | 70 | 70 | GUA | GUA | GUA | — | — | — | — | — | — | — | — | — |
| CO1 | + | 5481 | 7031 | 5480 | 7030 | 5487 | 7037 | 1551 | 1551 | 1551 | — | — | — | GTG | TAA | GTG | TAA | GTG | TAA | 516 | 516 | 516 |
| Ser (S) | - | 7032 | 7102 | 7031 | 7101 | 7038 | 7108 | 71 | 71 | 71 | UGA | UGA | UGA | — | — | — | — | — | — | — | — | — |
| Asp (D) | + | 7106 | 7178 | 7105 | 7177 | 7112 | 7184 | 73 | 73 | 73 | GUC | GUC | GUC | — | — | — | — | — | — | — | — | — |
| CO2 | + | 7183 | 7873 | 7188 | 7878 | 7193 | 7883 | 691 | 691 | 691 | — | — | — | ATG | T-- | ATG | T-- | ATG | T-- | 230 | 230 | 230 |
| Lys (K) | + | 7874 | 7947 | 7879 | 7952 | 7884 | 7957 | 74 | 74 | 74 | UUU | UUU | UUU | — | — | — | — | — | — | — | — | — |
| ATP8 | + | 7949 | 8116 | 7956 | 8123 | 7959 | 8126 | 168 | 168 | 168 | — | — | — | ATG | TAA | ATG | TAA | ATG | TAA | 55 | 55 | 55 |
| ATP6 | + | 8107 | 8789 | 8114 | 8796 | 8117 | 8799 | 683 | 683 | 683 | — | — | — | ATG | TA- | ATG | TA- | ATG | TA- | 227 | 227 | 227 |
| CO3 | + | 8790 | 9574 | 8797 | 9581 | 8800 | 9584 | 785 | 785 | 785 | — | — | — | ATG | TA- | ATG | TA- | ATG | TA- | 261 | 261 | 261 |
| Gly (G) | + | 9575 | 9646 | 9582 | 9653 | 9585 | 9655 | 72 | 72 | 71 | UCC | UCC | UCC | — | — | — | — | — | — | — | — | — |
| ND3 | + | 9647 | 9995 | 9654 | 10002 | 9656 | 10004 | 349 | 349 | 349 | — | — | — | ATG | T-- | ATG | T-- | ATG | T-- | 116 | 116 | 116 |
| Arg (R) | + | 9996 | 10064 | 10003 | 10071 | 10005 | 10073 | 69 | 69 | 69 | UCG | UCG | UCG | — | — | — | — | — | — | — | — | — |
| ND4L | + | 10065 | 10361 | 10072 | 10368 | 10074 | 10370 | 297 | 297 | 297 | — | — | — | ATG | TAA | ATG | TAA | ATG | TAA | 98 | 98 | 98 |
| ND4 | + | 10355 | 11735 | 10362 | 11742 | 10364 | 11744 | 1381 | 1381 | 1381 | — | — | — | ATG | T-- | ATG | T-- | ATG | T-- | 460 | 460 | 460 |
| His (H) | + | 11736 | 11804 | 11743 | 11812 | 11745 | 11813 | 69 | 70 | 69 | GUG | GUG | GUG | — | — | — | — | — | — | — | — | — |
| Ser (S) | + | 11805 | 11871 | 11813 | 11879 | 11814 | 11880 | 67 | 67 | 67 | GCU | GCU | GCU | — | — | — | — | — | — | — | — | — |
| Leu (L) | + | 11880 | 11952 | 11887 | 11959 | 11887 | 11959 | 73 | 73 | 73 | UAG | UAG | UAG | — | — | — | — | — | — | — | — | — |
| NCR | +/- | — | — | — | — | 11960 | 12179 | — | — | 220 | — | — | — | — | — | — | — | — | — | — | — | — |
| Leu (L) | + | — | — | — | — | 12180 | 12252 | — | — | 73 | — | — | UAG | — | — | — | — | — | — | — | — | — |
| ND5 | + | 11953 | 13791 | 11960 | 13798 | 12253 | 14091 | 1839 | 1839 | 1839 | — | — | — | ATG | TAG | ATG | AGA | ATG | TAA | 612 | 612 | 612 |
| ND6 | - | 13788 | 14309 | 13794 | 14315 | 14088 | 14609 | 522 | 522 | 522 | — | — | — | ATG | TAA | ATG | TAG | ATG | TAA | 173 | 173 | 173 |
| Glu (E) | - | 14310 | 14378 | 14316 | 14384 | 14610 | 14677 | 69 | 69 | 68 | UUC | UUC | UUC | — | — | — | — | — | — | — | — | — |
| CYTB | + | 14386 | 15526 | 14388 | 15528 | 14683 | 15823 | 1141 | 1141 | 1141 | — | — | — | ATG | T-- | ATG | T-- | ATG | T-- | 380 | 380 | 380 |
| Thr (T) | + | 15527 | 15598 | 15529 | 15600 | 15824 | 15895 | 72 | 72 | 72 | UGU | UGU | UGU | — | — | — | — | — | — | — | — | — |
| Pro (P) | - | 15599 | 15668 | 15601 | 15670 | 15896 | 15965 | 70 | 70 | 70 | UGG | UGG | UGG | — | — | — | — | — | — | — | — | — |
| Control region | + | 15669 | 16765 | 15671 | 16602 | 15696 | 16896 | 1097 | 932 | 1200 | — | — | — | — | — | — | — | — | — | — | — | — |

**Fig. 2.**
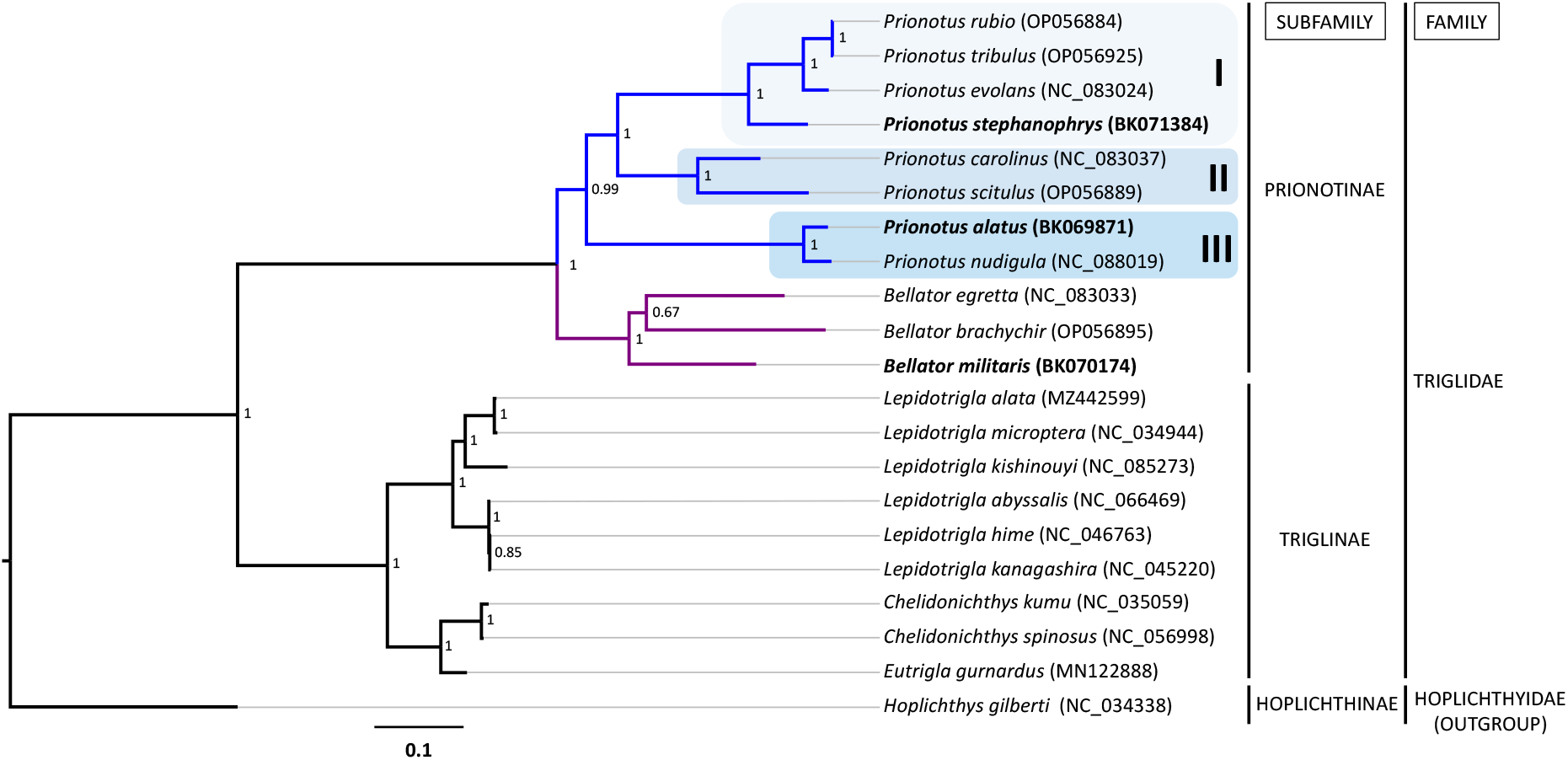
Bayesian phylogenetic tree constructed based on the concatenation of 13 mitochondrial protein-coding genes from 20 Triglidae species (GenBank accessions are provided in parenthesis). The three species with newly determined mitogenomes, *Bellator militaris, Prionotus alatus*, and *P. stephanophrys* (subfamily Prionotinae), are highlighted in bold. The nodes and branches from the *Bellator* and *Prionotus* groups are shown in violet and blue colors respectively. The three subclades formed within the *Prionotus* group are shaded in sky blue panels and denoted with Roman numbers (I, II, and III). Posterior probabilities are shown at the corresponding nodes. The Gilbert’s spiny flathead, *Hoplichthys gilberti*, was used as outgroup.

Although the unusual non-coding region observed in the four previously mentioned *Prionotus* species may have resulted from an unknown evolutionary process due to their close phylogenetic relationship (Fig. 2), we cannot entirely dismiss the possibility that it could have an artefactual origin. This is particularly relevant given the presence of a C homopolymer at the 5’ end of the non-coding region, which is known to introduce biases in short-read sequencing (Laehnemann et al. 2016).

The sizes of the PCGs ranged from 168 bp (55 amino acids) in the ATP8 gene to 1,839 bp (612 amino acids) in the ND5 gene from the three mitogenomes. All PCGs were encoded in the heavy strand and transcribed counterclockwise, except for the ND6 gene, which was located in the light strand and transcribed in a clockwise direction. Most PCGs initiated with the start codon ATG; however, the CO1 gene started with the alternative codon GTG in all mitogenomes. The use of alternative stop codons was observed only in the ND5 gene of the *B. militaris* mitogenome, which ended with the TAG codon. In the mitogenome of *P. alatus*, two genes, the ND5 and the ND6, ended with the AGA and TAG codons, respectively. In the mitogenome of *P. stephanophrys*, the ND1 gene also used the TAG stop codon (Table 2). In the three mitogenomes, the heavy strand contained the two rRNA and 14 tRNA genes, which included an additional tRNA-Leu found only in *P. stephanophrys*. The light strand contained the remaining 8 tRNA genes in all the reported mitogenomes.

### 3.3 Phylogenetic analysis

The Bayesian phylogenetic analysis provided strong to maximum statistical support for the clades Prionotinae and Triglinae (Fig. 2). In our phylogenetic tree, the subfamily Triglinae, which includes 9 species across 3 genera (*Chelidonichthys, Eutrigla*, and *Lepidotrigla*), was positioned basally within the family Triglinidae. This finding aligns with part of the molecular phylogeny proposed by Portnoy et al. (2017), which classified the Triglinae group as more basal than Prionotinae. However, this result contrasts with the morphological phylogenetic analysis conducted by Richards and Jones (2002), which identified the Prionotinae as the most ancestral subfamily within Triglidae.

All species within the genera *Bellator* and *Prionotus* were classified under a major monophyletic Prionotinae clade, supported by a maximum Bayesian posterior probability (BPP) value of 1. Notably, our results revealed that the *Bellator* group, which includes *B. egretta, B. brachychir*, and *B. militaris*, has a monophyletic origin and shows a well-supported sister group relationship with *Prionotus* (Fig. 2). This supports the current classification of *Bellator* as a distinct genus, in line with the morphological taxonomic studies conducted by Miller and Richards (1991), Imamura (1996), and Richards and Jones (2002). However, these results contrast with the partial gene-based phylogeny reported by Portnoy et al. (2017), which found *Prionotus* to be paraphyletic within Prionotinae and placed *Bellator* within the genus *Prionotus*.

The analysis of the eight *Prionotus* species revealed three distinct subclades, as illustrated in Fig. 2. These subclades are as follows: subclade I includes *P. rubio, P*. tribulus, *P. evolans*, and *P. stephanophrys*; subclade II comprises *P. carolinus* and *P. scitulus*; and subclade III consists of *P. alatus* and *P. nudigula*. Our findings indicate that *P. alatus* and *P. stephanophrys* are positioned in different subclades within the Prionotinae. Specifically, *P. alatus* has a closer evolutionary relationship with *P. nudigula* (subclade III), while *P. stephanophrys* is more closely related to *P. evolans* (subclade I). The estimated divergence levels (using the Kimura two-parameter model) between *P. alatus* and *P. nudigula* is 0.05, whereas the divergence between *P. stephanophrys* and *P. evolans* is 0.12. It is important to note that *P. stephanophrys* is the only Pacific species of *Prionotus* included in our study, and there are still several Atlantic species whose mitogenomes have yet to be sequenced. Therefore, we cannot identify which Atlantic species serves as the transisthmian sister taxon of *P. stephanophrys*. However, based on findings reported by Portnoy et al. (2017), it appears that *P. longispinosus* is the closest Atlantic relative of *P. stephanophrys* and likely represents its transisthmian sister species.

The shortest interspecific genetic distance (K2P=0.0014; 23 variable sites) was observed between the mitogenomes of *P. rubio* (Genbank accession OP056884, Biosample SAMN29555344) and *P. tribulus* (Genbank accession OP056925, BioSample SAMN29555068), both of which share the same length of 16,891 bp. A further analysis of the CO1 gene sequence from both mitogenomes in the BOLD database identified both sequences as *P. tribulus*. These findings suggest that Biosample SAMN29555344, which is identified as *P. rubio*, may be actually a hybrid of a male *P. rubio* and a female *P. tribulus*. Hybridization between different species of *Prionotus* has been reported both in the wild and under laboratory conditions (McClure and McEachran, 1992; Herbert et al. 2024), indicating that potential hybridization between the sympatric species *P. rubio* and *P. tribulus* is plausible.

## 4. Conclusion

We conducted the first complete mitogenome-based phylogenetic analysis of species within the Prionotinae subfamily. Our findings indicate a monophyletic origin for both sister taxa, *Bellator* and *Prionotus*, which aligns with some previous studies. However, this monophyletic relationship contradicts the conclusions of Portnoy et al. (2017), who suggested a paraphyletic origin for *Prionotus* and proposed that *Bellator* should be considered a junior synonym of *Prionotus*. It is worth noting that out of the three *Bellator* species included in both our analysis and that of Portnoy et al. (2017), *B. militaris* is the only *Bellator* species analyzed in both studies. Therefore, more extensive molecular phylogenetic analyses, incorporating a wider range of taxa and complete mitogenome sequences of additional *Bellator* and *Prionotus* species, are necessary to clarify the current genus status of *Bellator* and to validate the proposed monophyletic origin of *Prionotus*.

## Ethical approval

Not applicable.

## Informed consent

Not applicable.

## Data availability

Nucleotide sequence data reported are available in the Third Party Annotation Section of the DDBJ/ENA/GenBank databases under the accession number TPA: BK069871, BK070174, and BK071384.

